# Impact of exogenous nitrogen on cyanobacterial abundance and community in oil-contaminated sediment mesocosms

**DOI:** 10.1101/322743

**Authors:** Kaikai Sun, Minbo Qi, Panpan Zhu, Haixia Wang, Zhenmei Lu

## Abstract

High concentrations of crude oil are toxic to cyanobacteria but can facilitate the emergence of cyanobacterial aggregation at an appropriate concentration range; however, the exact inducing factor has never been clearly elucidated. We hypothesized that increasing exposure to elevated concentrations of nitrogen would inhibit the accumulation of cyanobacteria in oil-contaminated sediments. To test this hypothesis, we simulated an oil spill in estuarine sediment microcosms with and without the removal of nitrogen limitation by supplementation of exogenous nitrogen. An integrated MiSeq sequencing of 16S rRNA gene analysis along with metatranscriptome sequence was performed to achieve a comprehensive study of cyanobacterial blooms. The number of cyanobacterial sequences increased over time in both oil-contaminated and non-oil-contaminated sediments at different time points after 42 days of incubation. And, supplementation with a nitrogen resource could accelerate cyanobacterial blooms under uncontaminated microcosms but delay the bloom phenomenon and reduce the cyanobacterial abundance to a great degree when exposed to oil. Our results clearly illustrated that nitrogen limitation was a vital driver of the increased abundance of cyanobacteria in oil-contaminated mesocosms. In addition, the abundance and compositions of blooming cyanobacteria varied significantly among the different treatment groups, and *Oscillatoria* may play a potential and non-negligible role in oil-contaminated mesocosms.

## IMPORTANCE

Previous research has shown that cyanobacterial blooms do occur in oiled mesocosms and it is still unknown exactly the main driver for the induced cyanobacteria outbreaks. In this study, we investigated the abundance and composition of cyanobacteria in sediment mesocosms with or without supplemented exogenous nitrogen at DNA and RNA level. We revealed an interesting perspective that exogenous nitrogen played an opposite role of inducing cyanobacteria in the presence or absence of crude oil. Furthermore, it was possible to link previous phenomenon to changes of the abundance and compositions of blooming cyanobacteria. This work contributes novel knowledge on the population distributions and the mechanism of cyanobacterial bloom in oiled mesocosms. Moreover, the artificial addition of nitrogen could enhance crude oil degradation and delay accumulation of cyanobacterial harmful algal blooms (CyanoHABs), which might be one of the possible ways to ameliorate the threat of eutrophication in oil-contaminated sediments.

## Introduction

Cyanobacteria is considered significant contributors to the food chain; as a collective, cyanobacteria have the physiological capacity to occupy all aquatic environments irrespective of salinity: freshwater lakes, estuaries, marine oceans, and hypersaline mats (1). In general, crude oil is toxic to cyanobacteria, but it can stimulate cyanobacterial abundance at low concentrations in some cases (2). It has been reported that cyanobacteria can tolerate and develop in oil-contaminated regions (3, 4); cyanobacterial blooms do occur in oil-contaminated areas at certain concentrations (5, 6, 7). Some species of cyanobacteria are thought to participate directly in oil degradation as hydrocarbon degraders (8, 9). Additionally, cyanobacteria can generally perform indirect functions in the metabolism and degradation of different hydrocarbon compounds, for example, providing oxygen-fixing oil degraders to facilitate hydrocarbon metabolism (10, 11). However, a large-scale outbreak of cyanobacteria may cause secondary pollution. A large number of growing phytoplankton or cyanobacteria can seal the surface to produce an anaerobic environment, thus preventing microbial oil degradation. Moreover, some kinds of cyanobacteria produce a series of toxic cyanotoxins, such as microcystin, which harm aquatic animals and ecology systems. Death of cyanobacteria can also release a great quantity of hazardous substances and consume mass (dissolved oxygen).

The reasons underlying cyanobacterial blooms in oil-contaminated areas still remain unclear. Nonetheless, crude oil can inhibit the key deposit feeders, which is likely to be a major factor responsible for an increase in cyanobacteria abundance; furthermore, a low nitrogen concentration also presumably selects for cyanobacteria, especially dinitrogen-fixing cyanobacteria (6). N_2_ fixation is a key process in photosynthetic microbial mats to support nitrogen demands associated with primary production. In addition, certain cyanobacteria have the capacity to fix atmospheric dinitrogen (N_2_) and therefore significantly contribute to biogeochemical cycling of nitrogen in both aquatic and terrestrial ecosystems (12). Increased anthropogenic nutrient addition may alter the biogeochemical coupling between estuarine sediments and waters (13), which can further exacerbate eutrophic conditions (14), change the microbial ecology of the sediment, and in turn, accelerate cyanobacterial blooming process in marine or terrestrial systems. However, how anthropogenic nutrient functions in the whole bacterial or cyanobacterial community in oil-polluted areas when supplied with anthropogenic nitrogen still remains ambiguous.

We hypothesize that increasing exposure to an elevated concentration of nitrogen would inhibit the accumulation of cyanobacteria in oil-contaminated sediments. Hence, in this study, we simulated an oil spill in estuarine sediment microcosms with or without exogenous nitrogen supplementation and combined the MiSeq sequencing of 16S rRNA genes and metatranscriptome analysis to address the following questions: (i) What is the key induced factor of cyanobacterial blooms when exposed to crude oil? How do cyanobacteria function in oil-contaminated microcosms in the presence or absence of exogenous nitrogen? (iii) What are the differences in cyanobacterial(especially nitrogen-fixing cyanobacteria) composition among all the mesocosms?

## MATERIALS AND METHODS

### Sample collection and processing

Sediments were collected from the estuary of the Yong River (N29°57′56.35″, E121°44′6.91″; altitude, 5 m) on October 15, 2016. To recreate oil-contaminated conditions, a simulation system of microcosms was assembled with four containers that were all divided into three equal 20 × 10 × 30 cm (l × w × h) segments. Each reservoir contained 800 g fresh sediment topped with approximately 2 cm filtered seawater. Four treatments were as follows: a crude oil-contaminated microcosm with supplemental nitrogen group (CN), a crude oil-contaminated microcosm without supplemental nitrogen group (CO), an uncontaminated microcosm with supplemental nitrogen group (ON), and an uncontaminated microcosm without supplemental nitrogen group (OO). The crude oil concentration was 30 g kg^−1^ dry weight (wt.). Nitrogen concentrations were measured every 24 hours, and a ratio of four times exogenous nitrogen to that of the original sediment (OS) was added to reservoirs of the CN and ON groups when their depletion reached the relative lower value. All containers were placed in an illumination incubator with a temperature of 28 °C, relative humidity of 70%, and 3000 lux of illumination intensity. For geochemical and microbiological analyses, samples of the overlying water were collected every 24 hours; samples of sediments taken from the top 1.5 cm (days 0, 2, 5, 9, 12, 15, 19, 26, 33 and 42) were collected in triplicate: one was dried for sediment characteristic parameter analysis, one was stored at −20 °C in a freezer for DNA extraction, and one stored at −80 °C in a freezer for RNA extraction.

### Chemical analysis

To analyze ammonium nitrogen (NH_4_^+^-N), nitrate nitrogen (NO_3_^−^-N), nitrite nitrogen (NO_2_^−^-N) and the pH value of the overlying water (pre-filtered with 0.45-μm pore size filters), samples were taken to the laboratory and analyzed within 24 hours after sampling according to standard methods (EPAC, 2002).

The total nitrogen (TN), total phosphorus (TP) and total organic carbon (TOC) of sediments were measured by the Kjeldahl method, molybdenum blue method and spectrophotometric method with potassium dichromate, respectively.

### Nucleic acid extraction

DNA extractions were performed using a Power Soil DNA Isolation Kit (Mo Bio Laboratories, Carlsbad, CA, USA) according to the manufacturer’s instructions. An RNA PowerSoil^®^ Total RNA Isolation Kit was used for RNA extraction; 2 g of fresh samples were prepared for total RNA isolation from the organisms in the sediment. DNA and RNA quantity and quality were then measured with a Nanodrop spectrophotometer (Nanodrop ND1000, Wilmington, DE, USA) at a wavelength of 260 nm, and the integrity was evaluated via 1% of agarose gel electrophoresis. The RNA quality was also assessed by an Agilent 2100 Bioanalyzer (Agilent Technologies, Palo Alto, CA, USA).

### MiSeq analysis of the microbial community

Sequencing was performed on an Illumina MiSeq platform. Raw Illumina fastq files were demultiplexed, quality filtered and analyzed using QIIME (v. 1.4.0-dev) 7 (15) according to the standard protocols. Demultiplexed sequences were clustered into operational taxonomic units (OTUs) using the QIIME UCLUST12 wrapper, with a threshold of 97% pairwise nucleotide sequence identity, and the cluster centroid for each OTU was chosen as the OTU representative sequence, with taxonomy assigned to representative OTUs from each cluster using the Ribosomal Database Project classifier in QIIME, trained on the Greengenes database at 97% identity. After taxonomic assignment, variable OTU minimum abundance thresholds were applied to remove any OTU representing fewer sequences than the defined threshold. Representative sequences were aligned using PyNAST15 against a template alignment of the Greengenes database and filtered at 97% identity. To compare the community composition between samples, sequences were aligned using the PyNAST aligner in QIIME. This metric compares samples based on the phylogenetic relatedness (branch lengths) of OTUs in a community while taking into account the relative OTU abundance (16).

### Metatranscriptome library preparation and sequencing

One microgram of total RNA with an RIN value above 7 was used for the following library preparation. Next-generation sequencing library preparations were constructed according to the manufacturer’s protocol (NEBNext^®^ Ultra^TM^ Directional RNA Library Prep Kit for Illumina^®^). The rRNA was depleted from total RNA using a Ribo-Zero rRNA Removal Kit (Bacteria) (Illumina). The ribosomal-depleted mRNA was then fragmented and reverse-transcribed. First-strand cDNA was synthesized using ProtoScript II Reverse Transcriptase with random primers and actinomycin D. The second-strand cDNA was synthesized using Second Strand Synthesis Enzyme Mix (including dACG-TP/dUTP). The double-stranded cDNA purified by AxyPrep Mag PCR Clean-up (Axygen) was then treated with End Prep Enzyme Mix to repair both ends and to add a dA-tailing in one reaction, followed by T-A ligation to add adaptors to both ends. Size selection of adaptor-ligated DNA was then performed using AxyPrep Mag PCR Clean-up (Axygen), and fragments of ~360 bp (with an approximate insert size of 300 bp) were recovered. The dUTP-marked second strand was digested with the Uracil-Specific Excision Reagent (USER) enzyme (New England Biolabs). Each sample was then amplified by PCR for 11 cycles using P5 and P7 primers, with both primers carrying sequences that can anneal with flow cells to perform bridge PCR and a P7 primer carrying a six-base index allowing for multiplexing. The PCR products were cleaned up using AxyPrep Mag PCR Clean-up (Axygen), validated using an Agilent 2100 Bioanalyzer (Agilent Technologies, Palo Alto, CA, USA), and quantified by a Qubit 2.0 Fluorometer (Invitrogen, Carlsbad, CA, USA). Then, libraries with different indices were multiplexed and loaded onto an Illumina HiSeq instrument according to the manufacturer’s instructions (Illumina, San Diego, CA, USA). Sequencing was carried out using a 2×150 paired-end (PE) configuration; image analysis and base calling were conducted by the HiSeq Control Software (HCS) + OLB + GAPipeline-1.6 (Illumina) on the HiSeq instrument.

Raw shotgun sequencing reads were trimmed using cutadapt (v1.9.1) Low-quality reads, Nrich reads and adapter-polluted reads were removed. Then host contamination reads were removed. rRNA reads were repeatedly removed using SortMeRNA aligning to the SILVA 128 version database. The PE reads were assembled using Trinity. Trinity uses the de *Bruijn* graph strategy to assemble the transcriptome. Open reading frames (ORFs) were identified using TransDecoder program, with default parameters. All sequences with a 95 % sequence identity (90% coverage) were clustered as the non-redundant gene catalog by the CD-HIT. BLASTP (Version 2.2.28+) was employed for taxonomic annotations by aligning non-redundant gene catalogs against the NCBI nonredundant (NR) protein database with e-value cutoff of 1e-5. Reads after quality control were mapped to the representative genes with 95% identity and FPKM were evaluated using RSEM.

### Quantitative analysis of cyanobacterial 16S rRNA genes

PCR amplifications were performed using Rotor-gene Q (QIAGEN, Germany) and carried out in a reaction mixture containing 10 μL of 1× SYBR Premix Ex Taq, 1 μL of forward and reverse primers (10 μM), and 8 μL of DNA template. The DNA template was diluted with sterilized distilled water to make sure that approximately 20 ng of DNA was included in each reaction mixture. DNA was obtained from the core on days 0 and 26 (previously used for 16S rRNA gene pyrosequencing analysis).

Quantitative real-time PCR (qPCR) analysis of cyanobacteria was carried out using the cyanobacteria-specific primer set Cya359/Cya781A-B (17), and the V6 region of the 16S rRNA gene, amplified with primers 984F/1378R, was selected as the reference gene. The qPCR cycling conditions were as follows: initial denaturation at 95°C for 3 min, followed by 40 cycles of denaturation at 95 °C for 10 s, annealing at 57 °C for 10 s and extension at 72 °C for 6 s.

### Clone libraries of the *nifH* gene

Universal primers for *nifH* (18) were used for amplification of DNA, with an initial denaturation at 94°C for 1 min; followed by 40 cycles of 1 min denaturation at 94°C, 2 min annealing at 57°C and 2 min extension at 72°C; with a final step of extension for 5 min. For samples that produced the appropriately sized product, the band was excised and purified with a QIAquick Gel Purification Kit (Qiagen) according to the manufacturer’s protocol. Purified PCR products were eluted in 35 μl of EB buffer, and 2 μl of the purified product was immediately ligated into the pGEM-T Easy Vector system (Promega, Madison, WI, USA) and transformed into chemically competent *Escherichia coli* cells following the manufacturer’s instructions. Cells, approximately 100 positive colonies per ligation, were inoculated into 1 ml of LB buffer overnight at 37°C, and plasmids were purified using a Montáge Plasmid Miniprep96 Kit (Millipore). All sequencing was performed on Applied Biosystems 48 capillary 3730 or Applied Biosystems 16 capillary 3100 Genetic Analyzer Systems in one direction with the M13F or M13R primer.

## Results

### Environmental parameters and degradation of total petroleum hydrocarbons

The variation in NH_4_^+^-N, NO_2_^−^-N, NO_3_^−^-N in the overlying water was detected during the operation period (Fig. 1). When nitrogen in the system was almost completely consumed resulted from rapid depletion of NH_4_^+^-N, NO_2_^−^-N, NO_3_^−^-N during the initial five days, additional nitrogen was supplemented to mudflat sediment mesocosms of the CN and ON groups to relieve nitrogen limitation with NH_4_Cl, NaNO_3_, and NaNO_2_ at day 6. After this supplementation, the nitrogen maintained an overall descending trend until it was exhausted. Ammonia consumption in the oil-contaminated groups lagged slightly behind that of the non-oil-contaminated groups, while nitrate and nitrite exhibited opposite results, indicating much greater nitrate and nitrite demand in the oil-contaminated sediments.

**Fig. 1.**
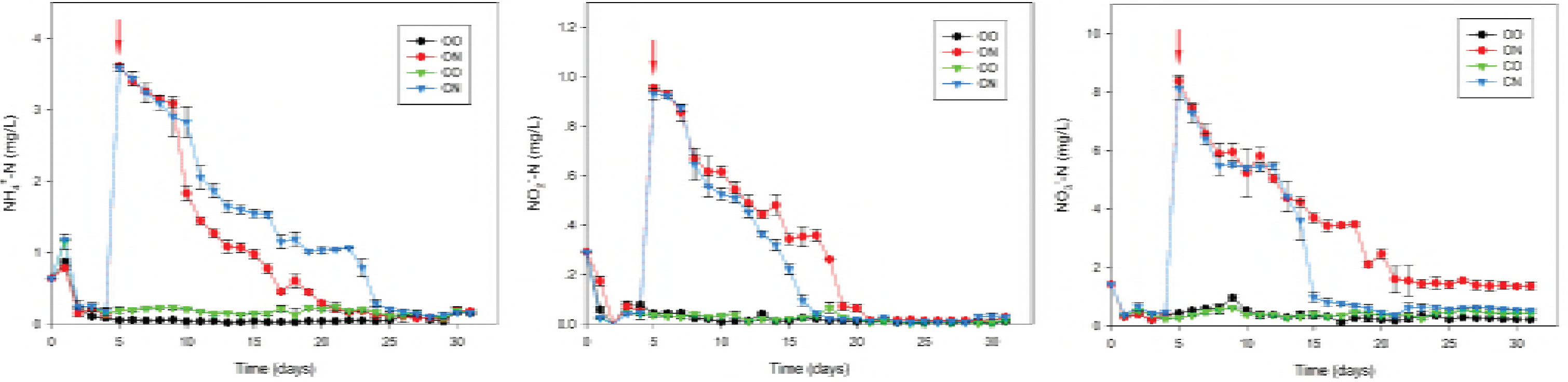
Changes in the physical and chemical properties in the overlying water column for the different groups. Error bars represent standard deviations based on analysis of three replicates for each sample. The red arrows indicate supplementation with exogenous nitrogen. OO, the uncontaminated microcosm without supplemental nitrogen group; ON, the uncontaminated microcosm with supplemental nitrogen group; CO, the crude oil-contaminated microcosm without supplemental nitrogen group; CN, the crude oil-contaminated microcosm with supplemental nitrogen group.

As for the remaining total petroleum hydrocarbon (TPH) relative to day 0, rapid degradation was shown in the first 19 days in the oil-contaminated sediments, while the degradation of the CO group lagged significantly behind that of the CN group (Fig. S1). In addition, the percentage of TPH remaining in the CO group was 58.76%, while in the CN group, it was 45.11%, indicating an enhanced degradation capacity after nitrogen supplementation in the oil-contaminated sediments.

### Cyanobacteria bloomed at different times with unequal abundance among all mesocosms

Green mats appeared upon the sediment surface of microcosms at different time points among all treatments during 42 days of incubation. MiSeq analysis of 16S rRNA gene indicated that the number of cyanobacterial sequences increased over time with significant differences in both oil-contaminated and non-oil-contaminated sediments, which may further confirmed that cyanobacteria bloomed on the sediment surfaces (Fig. 2). The relative abundances of cyanobacteria did not change significantly during the initial 9 days, following notably increased relative abundances of cyanobacteria occurred firstly in the non-oil-contaminated microcosms, with samples from the ON group earlier than those from the OO group (on day 12 and day 15, respectively). However, the oil-contaminated treatments exhibited different corresponding pathways to the increased concentration of nitrogen. The relative abundance of cyanobacteria dramatically increased in the CO group after 19 days of incubation and subsequently in CN group after 26 days. Overall, the number of cyanobacterial sequences in the non-oil-contaminated sediments was 1.9 times higher than that in the oil-contaminated sediments; and 4.6 times higher in the nitrogen-addition sediments than that in the non-addition sediments meanwhile. For oil-contaminated sediments, the cyanobacterial sequences of the CO group were more abundant than those of the CN group. It is noteworthy that the relative abundance all increased in varying degrees except for the CN group after 42 days of incubation. Cyanobacterial abundance at the transcriptional level showed a similar result, where cyanobacteria comprised 33.58% of the sequencing fragments in the ON group on day 26, with 4.14%, 3.63%, 9.00% and 1.56% in the OS, OO, CO and CN groups, respectively (Fig. S2).

**Fig. 2.**
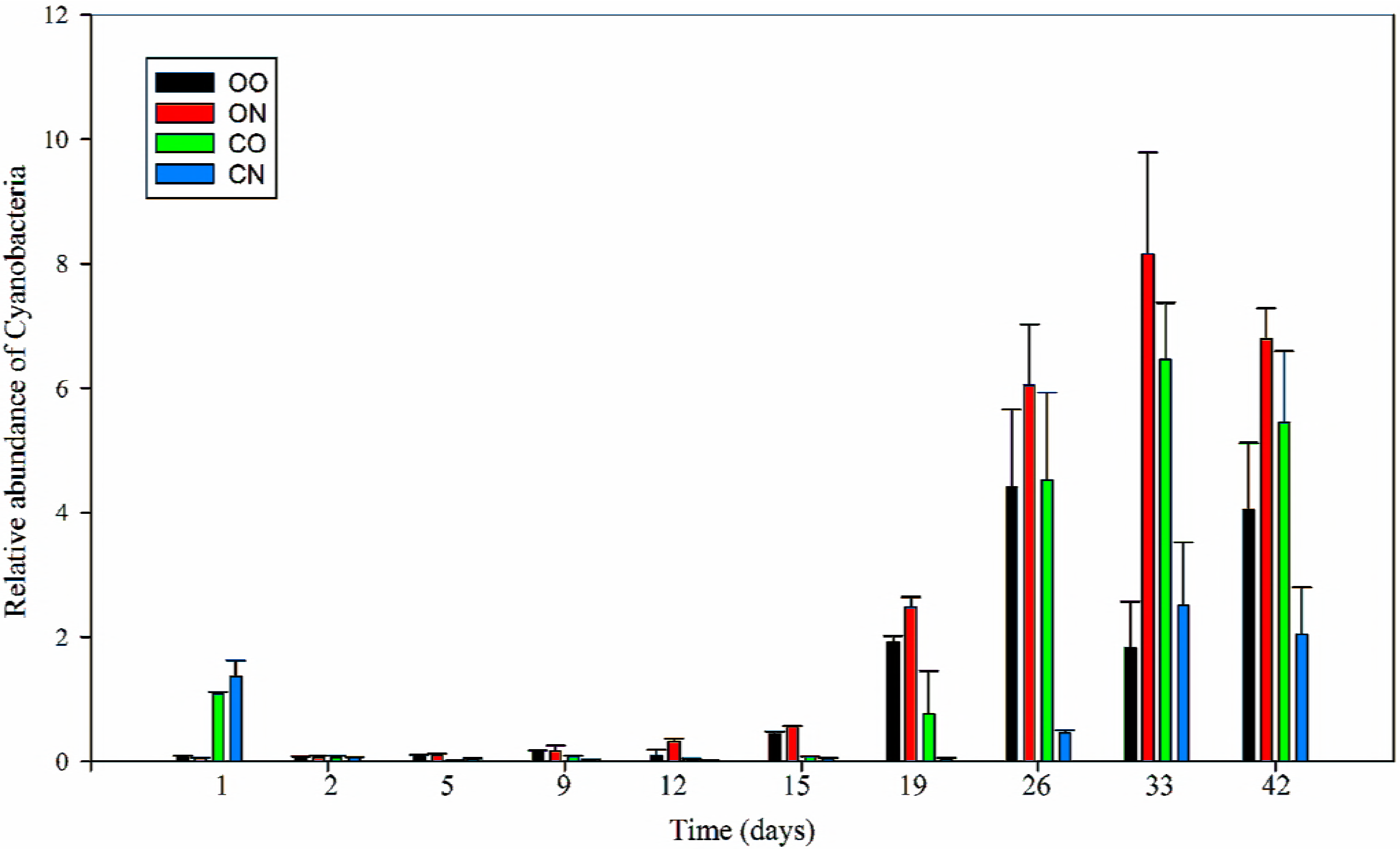
The relative abundance of cyanobacteria in the different groups as determined by MiSeq sequencing. Error bars represent standard deviations based on analysis of three replicates for each sample.

qPCR was also performed to quantitatively analyze cyanobacterial 16S rRNA genes using their specific primer pair (Fig. S3), and these genes exhibited trends similar to that of phylotype cyanobacterial abundance among all treatment microcosms. The relative abundance of cyanobacterial 16S rRNA genes did not change significantly during the initial incubation period, corresponding to the low abundance of Cyanobacteria from sequencing data. Furthermore, increased relative abundance of cyanobacterial 16S rRNA genes also occurred earlier in non-oil-contaminated microcosms than that in the oil-contaminated microcosms and in samples from the ON group earlier than those from the OO group. However, it seems that cyanobacterial 16S rRNA genes from the oil-contaminated treatments did not show a significant difference in response to an increased concentration of nitrogen. After 42 days of incubation, the cyanobacterial 16S rRNA gene levels all rose rapidly.

### Detection of nitrogenase genes and phylotypes

Genes coding for nitrogen fixation, such as *nifH*, were well represented in oiled metatranscriptomes, and similar activity was observed in oil-free control microcosms but not in the ON group (Fig. S4). Moreover, the nitrogen-fixation genes in the CO group were far more abundant than those in the other groups. To further verify this finding, a total of 439 clones were obtained from the combined *nifH* clone libraries from the original sediment and sediments on day 26 (Fig. 3), and most of them could be classified using BLASTN. Clone libraries of *nifH* genes revealed that the compositions of nitrogen-fixing bacteria on day 26 were obviously different from those of samples from the original sediment, indicating that crude oil and exogenous nitrogen do radically have a remarkable effect on the diversity of *nifH*-containing microbes. The most notably differences were sequences from cyanobacteria and *Marinobacterium lutimaris*, which were only detected in oil-contaminated mesocosms. Furthermore, the relative proportion of cyanobacteria in the CO group was obviously higher than that in the CN group.

**Fig. 3.**
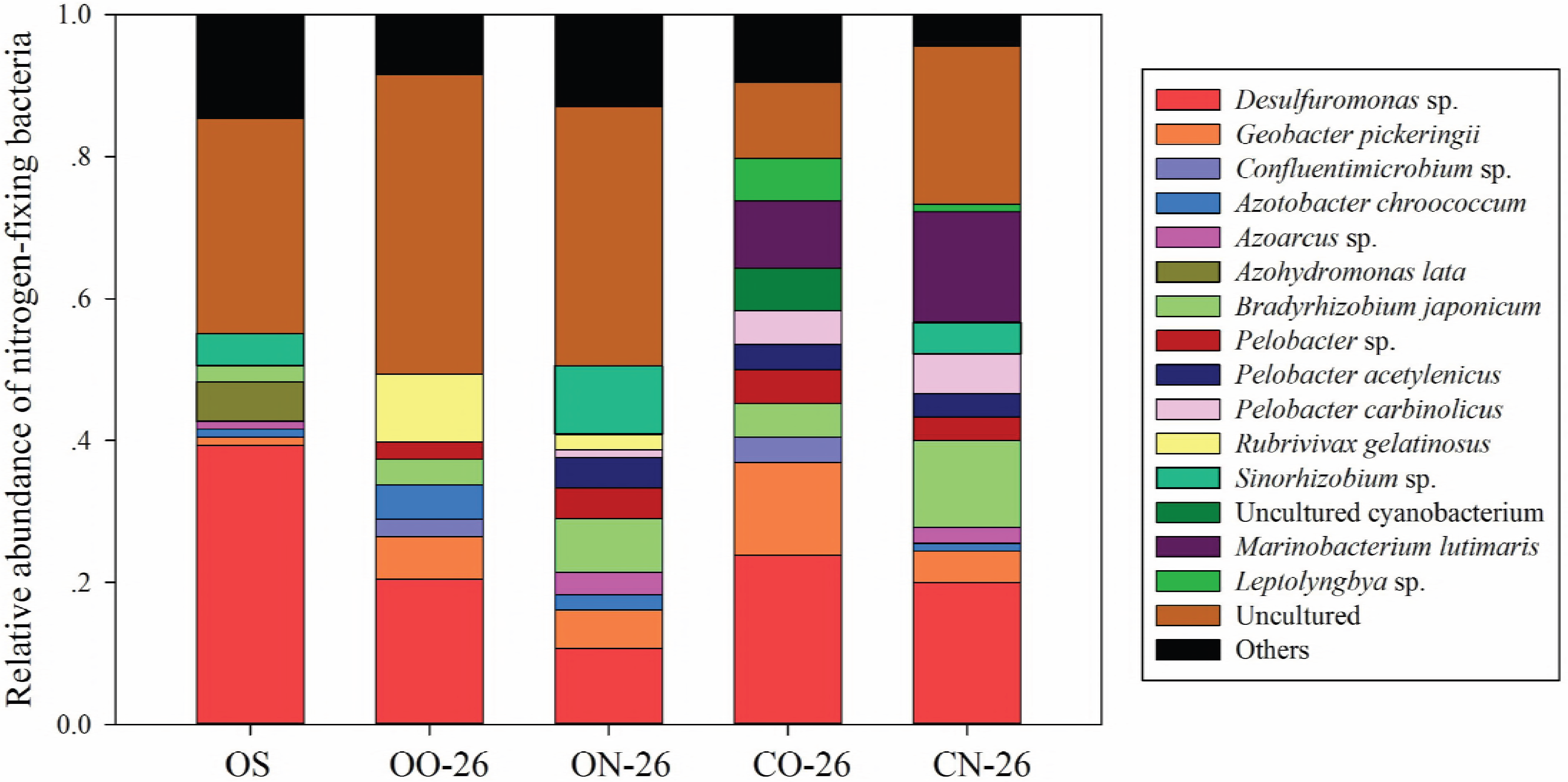
The relative abundance of nitrogen-fixing bacteria in the original sediment and on day 26. Less-abundant phyla are grouped under the category ‘others.’ OS were derived from the original sediment. OO-26, ON-26, CO-26, and CN-26 were derived from the uncontaminated microcosm without supplemental nitrogen group, the uncontaminated microcosm with supplemental nitrogen group, the crude oil-contaminated microcosm without supplemental nitrogen group, and the crude oil-contaminated microcosm with supplemental nitrogen group on day 26, respectively.

### The comprehensive cyanobacterial species composition of bloom-forming cyanobacteria

An examination of the cyanobacterial sequences revealed differences in the transcriptionally active cyanobacterial community structures among mesocosms (Fig. 4). At the order level, Oscillatoriales (~80%) holds an absolutely dominant position in the original sediment, in which most of the sequences were assigned to *Coleofasciculus chthonoplastes*. Apart from the original sediment, genera *Synechococcus, Oscillatoria* and *Leptolyngbya* were the most abundant taxa and were present at all four sampling mesocosms. The abundance of *Coleofasciculus chthonoplastes* dropped to a low level over time without any extra substance addition, and its predominance was replaced by that of *Synechococcus* and *Leptolyngbya. Oscillatoria* and *Synechococcus* were the dominant taxa, comprising 38.2% and 23.8% of the total cyanobacteria, respectively, after nitrogen supplementation. In addition, the relative abundance of *Leptolyngbya* was at a relative lower percentage (approximately 0.9%) in ON mesocosms compared with 24.5% in OO mesocosms, and it was more abundant in the non-oil-contaminated mesocosms than that in oil-contaminated mesocosms. Strains of *Synechococcus* dominated in the ON group and OO group, which also responded negatively to crude oil; besides, the strains of *Oscillatoria* showed a different distribution pattern. In our study, *Oscillatoria* occupied an absolute dominant position in oil-contaminated mesocosms, and the relative abundance of *Oscillatoria* in CO mesocosms was almost double that in CN mesocosms.

**Fig. 4.**
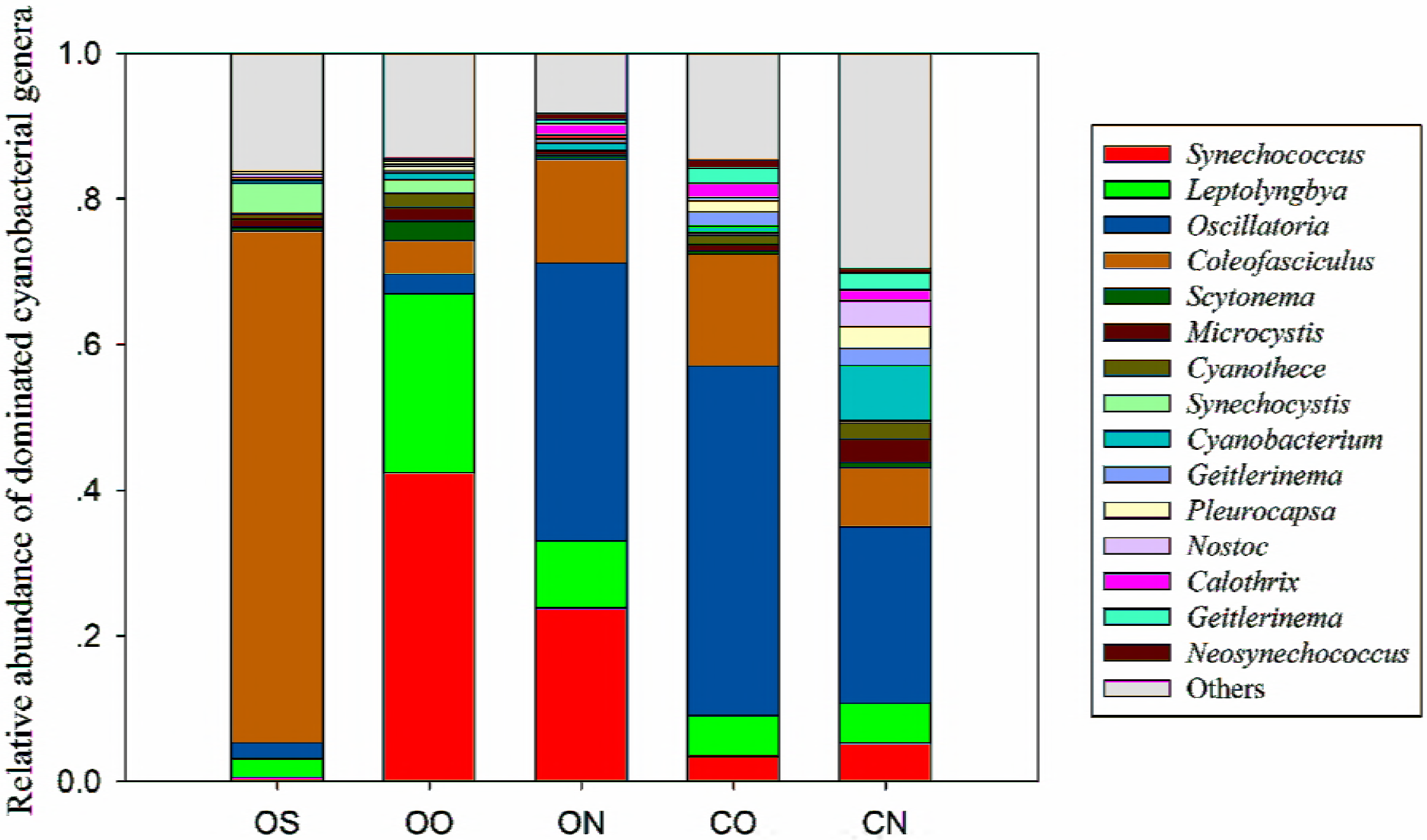
The relative abundances of the dominant cyanobacteria genera in different groups as determined by metatranscriptome sequencing. Less-abundant phyla are grouped under the category ‘others’.

Redundancy analysis (RDA) was applied to evaluate the potential relationship between environmental variables and cyanobacterial species (Fig. 5). *Synechococcus*, *Synechocystis*, *Scytonema*, *Leptolyngbya*, *Cyanothece*, and *Microcystis* correlated positively with NH_4_^+^-N and NO_2_^−^-N but negatively with pH. *Nostoc*, *Pleurocapsa*, *Pseudanabaena*, *Cyanothece*, *Neosynechococcus*, *Geitlerinema*, *Planktothricoides*, *Oscillatoria* and *Calothrix* correlated positively with TN, TP, TOC and NO_3_^−^-N but negatively with NH_4_^+^-N and NO_2_^−^-N. The value of environmental variables was listed in the Table S1.

**Fig. 5.**
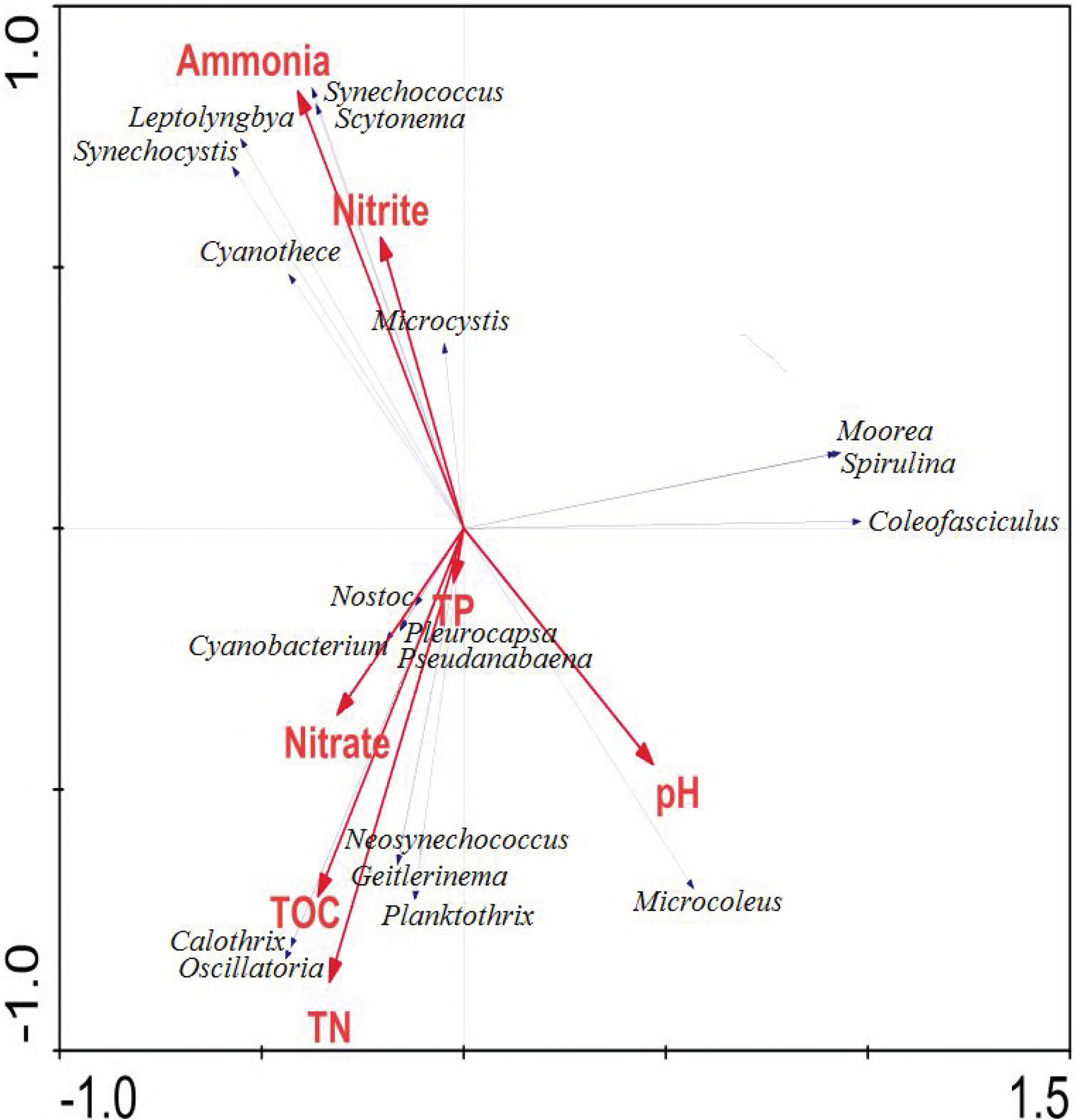
RDA ordination diagram of the data, with environmental variables represented by red arrows and cyanobacterial genera abundance represented by gray arrows. Correlations between environmental variables and RDA axes are represented by the length and angle of the arrows.

## Discussion

### Exogenous nitrogen delayed cyanobacterial blooms in oiled mesocosms

When nitrogen resources decreased to a very low concentration, cyanobacterial mats appeared on surfaces of both oil-contaminated and non-oil-contaminated sediments. Suitable growth conditions, with intense light, optimum temperature and appropriate humidity, can offer cyanobacteria a relative nutrient-rich environment in non-oil-contaminated microcosms to spur them to bloom (19). In addition, nitrogen supplementation yielded increasing aggravation of water eutrophication in the non-oil-contaminated systems and then induced a more rapid outbreak of cyanobacteria in the ON group compared to the OO group. Furthermore, many studies have proven that high concentrations of crude oil are toxic to cyanobacteria (20) but can facilitate the emergence of cyanobacterial aggregation at an appropriate concentration threshold (5, 6, 7, 21). Additional exogenous nitrogen could accelerate cyanobacterial blooms under uncontaminated microcosms; however, it did not play a similar function in the oil-contaminated systems but inhibited and delayed the enrichment of cyanobacteria no matter at DNA or RNA level in our study. We further explored the relative activity of photosynthesis and photosynthesis-antenna proteins involved in photosystem II operating efficiency (Fig. S5), since cyanobacteria are autotrophic prokaryotes that are capable of oxygenic photosynthesis similar to that of higher plants. On the one hand, it was clear that oil had an inhibitory effect on PS II efficiency at the late stage. Similarly, Piehler et al (22) and Chronopoulou et al (6) also found a reduction in photosynthetic efficiency in samples exposed to oil. On the other hand, nitrogen addition in oil-contaminated microcosms would further exacerbate the inhibitory effect of crude oil on PS II efficiency, which was corresponding to the relative abundance of cyanobacteria to some degree and further confirmed our conclusion.

### Reasons for changes in the relative abundance of cyanobacteria caused by exogenous nitrogen

Certain cyanobacteria have the capacity to fix atmospheric dinitrogen (N^2^) and therefore significantly contribute to the biogeochemical cycling of nitrogen in both aquatic and terrestrial ecosystems (12). In addition, Musat et al (23) demonstrated that cyanobacteria are the most active dinitrogen fixers under oil-contaminated conditions when nitrogen is limited. In some cases, the restriction of nitrogen can be eliminated or can undergo remission by dinitrogen fixation in the process of petroleum hydrocarbon degradation (24, 25). According to our results, artificially removing the nitrogen limitation of oil-contaminated microcosms delayed cyanobacterial blooming, which can confirm the standpoint that nitrogen limitation was a possible driver of the increased abundance of nitrogen-fixing cyanobacteria and changes in the cyanobacterial composition (6). To further verify this finding, genes involved in nitrogen fixation, such as *nifH*, which catalyzes dinitrogen fixation to ammonia, were analyzed. The result indicated that only the microcosms established under oil contamination without nitrogen supplement exhibited significant nitrogenase activity compared to others at day 26 (Fig. S3). It seems that oil was shown to have a driving role in dinitrogen fixation in oil-contaminated sediment microcosms in this study.

To our surprise, sequences related to *nifH* from cyanobacteria were only amplified from the oil-contaminated microcosms and constituted 11.9% and 5.56% of the CO and CN *nifH* library, respectively (Fig. 3). In addition, the cyanobacterial sequences obtained in these two *nifH* libraries were closely related to those of phylotypes *Leptolyngbya*, which were reported to be diazotrophs in ecological environments (26), and uncultured cyanobacteria. Nevertheless, even though the remaining oil-free samples had more cyanobacterial 16S rRNA gene sequences than the oil-contaminated samples, there was no cyanobacterial *nifH* sequence was detected among them. Therefore, cyanobacteria are the key oil-induced dinitrogen fixers in the oil-contaminated sediments but may play other ecological functions instead of nitrogen fixation in the non-oil-contaminated microcosms. In conclusion, nutrient limitation is a vital factor for stimulating cyanobacterial blooms in the oil-contaminated microcosms but is not a potential factor inducing cyanobacteria blooming in the non-oil-contaminated microcosms.

Many oil-degrading bacteria have been shown to carry out dinitrogen fixation (27, 28). Only relatively fewer bacterial species, *Desulfuromonas* and *Marinobacterium lutimaris*, reported to be capable of nitrogen fixation as well as oil degradation, were detected in *nifH* libraries (Fig. 3). However, it seems that the nitrogen-fixing capacity of hydrocarbon-degrading bacteria was not expressed during growth with hydrocarbons but only with polar substrates (29). In addition, several species of cyanobacteria were reported to show a high tolerance to oil and could participate directly in oil degradation as hydrocarbon degraders (8, 9, 27, 30). Therefore, the increased abundance of cyanobacteria may result from the need for hydrocarbon degraders in oil-contaminated mesocosms. However, the relative abundance of cyanobacteria increased at a low speed when rapid hydrocarbon degradation was exhibited during the initial 19 days (Fig. S1). Thus, a correlation between cyanobacterial abundance and hydrocarbon degradation was not observed. Many other studies have revealed that cyanobacteria themselves apparently could not degrade petroleum compounds directly but were more likely to play a significant, indirect role in biodegradation by supporting the growth and activity of symbiotic degraders (11, 31). In conclusion, the demand for hydrocarbon degradation was distinctly possible to be a minor factor for cyanobacterial blooming in oil-contaminated mesocosms.

### Comparison of cyanobacterial communities

Previous studies have estimated that populations and communities in the environment might be directly and/or indirectly affected by exposure to crude oil (22, 32, 33, 34). In general, nutrients are thought to be the most important environmental factor influencing the growth of phytoplankton. As above, both crude oil and nitrogen can influence cyanobacterial abundance, but currently, there is little understanding of their potential ability to regulate changes in cyanobacterial community composition. Our previous study showed that the phylotypes *Leptolyngbya*, *Oscillatoria*, *Arthrospira* (*Spirulina*), *Geitlerinema* and *Cyanothece* were dominant in the oil-contaminated sediments (21). In this study, the genera *Synechococcus*, *Oscillatoria* and *Leptolyngbya* were the most abundant taxa with high abundance, and these genera were present in all sampling mesocosms in the presence and absence of oil according to metatranscriptome data (Fig. 4). Strains of *Synechococcus*, a globally highly abundant genus, learned to survive in the oil-free groups. However, *Oscillatoria*, typical diazotrophs in ecological environments (26), occupied an absolutely dominant position in oil-contaminated mesocosms, suggesting that *Oscillatoria* may possess high tolerance to oil and play a potential and non-negligible role in oil-contaminated mesocosms. And the relative abundances of *Leptolyngbya* in the oil-contaminated mesocosms were lower than those in the non-oil-contaminated mesocosms, indicating that it may have a preference for a more suitable environment and, thus, likely plays other integral roles. The RDA ordination diagram (Fig. 5) clearly showed that the cyanobacterial community experienced rapid and substantial changes during the study. These changes are statistically related to various water and sediment quality variables, including NH_4_^+^-N, TN, TOC and pH. In this study, *Oscillatoria* demonstrated a stronger positive correlation with TN but a negative correlation with NH_4_^+^-N, while *Synechococcus* and *Leptolyngbya* showed the opposite result, suggesting that nitrogen supplementation, especially NH_4_^+^-N, did change the composition of cyanobacteria.

Excessive cyanobacterial (especially harmful algae) abundance has harmful effects on lake ecosystems and water quality, leading to the development of surface blooms and scums, oxygen depletion, and production of highly toxic cyanotoxins, which are extremely harmful to the environment and human health (35). In addition to being important primary producers, providing fixed carbon and nitrogen to higher trophic levels in the ecosystem (36, 37, 38), cyanobacteria can also produce a series of secondary metabolites, some with documented toxicity, such as cyanotoxins (39). In this study, cyanobacterial harmful algal blooms (CyanoHABs), such as typical *Anabaena*, *Planktothrix* and *Microcystis*, exhibited a pattern similar to that of other cyanobacteria, reducing their abundance in response to anthropogenic over-enrichment nitrogen in oil-contaminated sediments (Fig. S6). In part, this relationship was based upon the fact that some common CyanoHAB genera can fix atmospheric N_2_ into biologically available NH_3_, which could support the nitrogen requirements of blooming populations (40). Therefore, we may conclude that exogenous nitrogen can delay the accumulation of CyanoHABs and prevent secondary pollution under oil spills, to some extent.

## Conclusions

We observed large increases in cyanobacterial abundance among all treatment mesocosms but with different distributions between oil-contaminated and non-oil-contaminated mesocosms. Our results indicated that supplementation with a nitrogen resource could accelerate cyanobacterial blooms under non-oil-contaminated microcosms but delay bloom phenomenon and reduce abundance to a great degree when exposed to oil. Nitrogen limitation was an essential driver for the increased abundance of cyanobacteria in oil-contaminated areas. In addition, the anthropochory of extra nitrogen can enhance crude oil degradation capacity and delay the accumulation of CyanoHABs. Therefore, increased exogenous nitrogen input is one of the possible ways to ameliorate the threat of eutrophication in oil-contaminated sediments.

## ACKNOWLEDGMENTS

This work was financially supported by the National Natural Science Foundation of China (No. 31370151 and 31422003), and the Natural Science Foundation for Distinguished Young Scholars of Zhejiang Province (LR14D030001).

